# Global transcriptome analysis of *Aedes aegypti* mosquitoes in response to Zika virus infection

**DOI:** 10.1101/179416

**Authors:** Kayvan Etebari, Shivanand Hegde, Miguel A Saldaña, Steven G Widen, Thomas G Wood, Sassan Asgari, Grant L Hughes

**Author notes:** Corresponding authors: Sassan Asgari –. Grant L Hughes.

## Abstract

Zika virus (ZIKV) of the *Flaviviridae* family is a recently emerged mosquito-borne virus that has been implicated in the surge of the number of microcephaly instances in South America. The recent rapid spread of the virus led to its declaration as a global health emergency by the World Health Organization. The virus is transmitted mainly by the mosquito *Aedes aegypti* that also vectors dengue virus, however little is known about the interactions of the virus with the mosquito vector. In this study, we investigated the transcriptome profiles of whole *Ae. aegypti* mosquitoes in response to ZIKV infection at 2, 7, and 14 days post-infection using RNA-Seq. Results showed changes in the abundance of a large number of transcripts at each time point following infection, with 18 transcripts commonly changed among the three time points. Gene ontology analysis revealed that most of the altered genes are involved in metabolic process, cellular process and proteolysis. In addition, 486 long intergenic non-coding RNAs were identified that were altered upon ZIKV infection. Further, we found correlational changes of a number of potential mRNA target genes with that of altered host microRNAs. The outcomes provide a basic understanding of *Ae. aegypti* responses to ZIKV and helps to determine host factors involved in replication or mosquito host anti-viral response against the virus.

**Importance:** Vector-borne viruses pose great risks on human health. Zika virus has recently emerged as a global threat, rapidly expanding its distribution. Understanding the interactions of the virus with mosquito vectors at the molecular level is vital for devising new approaches in inhibiting virus transmission. In this study, we embarked on analyzing the transcriptional response of *Aedes aegypti* mosquitoes to Zika virus infection. Results showed large changes both in coding and long non-coding RNAs. Analysis of these genes showed similarities with other flaviviruses, including dengue virus, which is transmitted by the same mosquito vector. The outcomes provide a global picture of changes in the mosquito vector in response to Zika virus infection.

## Introduction

Flaviviruses are a group of arthropod-borne viruses (arboviruses) that impose huge burdens on global animal and human health. The most known examples of flaviviruses that cause diseases in humans are yellow fever, West Nile, dengue and Zika viruses. Zika virus (ZIKV) has been the most recently emerged mosquito-borne virus. While it was first reported in 1952 from Uganda (1), the virus spread rapidly across the Pacific and the Americas in the last 10 years with recent outbreaks in South America (2). The clinical symptoms are variable, ranging from no or mild symptoms to severe neurological disorders such as microcephaly in infants born from infected mothers, or Guillain-Barré syndrome in adults (reviewed in (2, 3)). The virus is mainly transmitted among humans by the bites of mosquito species of the genus *Aedes*, in particular *Aedes aegypti*, when they take a blood meal from infected individuals. The virus first infects the midgut cells of the mosquito and then disseminates into other tissues, finally reaching the salivary glands where they continue to replicate and are eventually transmitted to other human hosts upon subsequent blood feeding events (4).

It is thought that infection by flaviviruses does not cause any detrimental pathological effects on the mosquito vectors (5), reflecting evolutionary adaptations of the viruses with mosquitoes through intricate interactions, which involve optimal utilization of host factors for replication and avoidance of overt antiviral responses. However, a number of studies have shown major transcriptomic changes in the mosquito vectors in response to flavivirus infection. These changes suggest regulation of a wide range of host genes involved in classical immune pathways, RNA interference, metabolism, energy production and transport (6-13). In addition, mosquito small and long non-coding RNAs have also been shown to change upon flavivirus infection (14, 15).

Recently, we showed that the microRNA (miRNA) profile of *Ae. aegypti* mosquitoes is altered upon ZIKV infection at different time points following infection (16). Here, we describe the transcriptional response of *Ae. aegypti* whole mosquitoes to ZIKV infection at the same time points post-infection. Consistent with previous studies on other arboviruses, we found that the abundance of a large number of genes was altered following ZIKV infection.

## Results and Discussion

### *Ae. aegypti* RNA-Seq data analysis

RNA sequencing using Illumina sequencing technology was performed on poly(A)- enriched RNAs extracted from ZIKV-infected and non-infected *Ae. aegypti* mosquitoes at 2, 7, and 14 days post-infection (dpi). Total numbers of clean paired reads varied between 43,486,502 to 60,486,566 reads per library among the 18 sequenced RNA samples. More than 96% of reads mapped to the host genome with around 80% of counted fragments mapped to gene regions and 20% to intergenic areas of the genome (Table S1).

Principal component analysis (PCA) of the RNA-Seq data at each time point distributed all biological replicates of ZIKV-infected and non-infected samples in two distinct groups, although the differences were more subtle at 2 days post-infection, in which one of the ZIKV-infected biological replicates was relatively close to the control group (Fig. 1).

**Figure 1.**
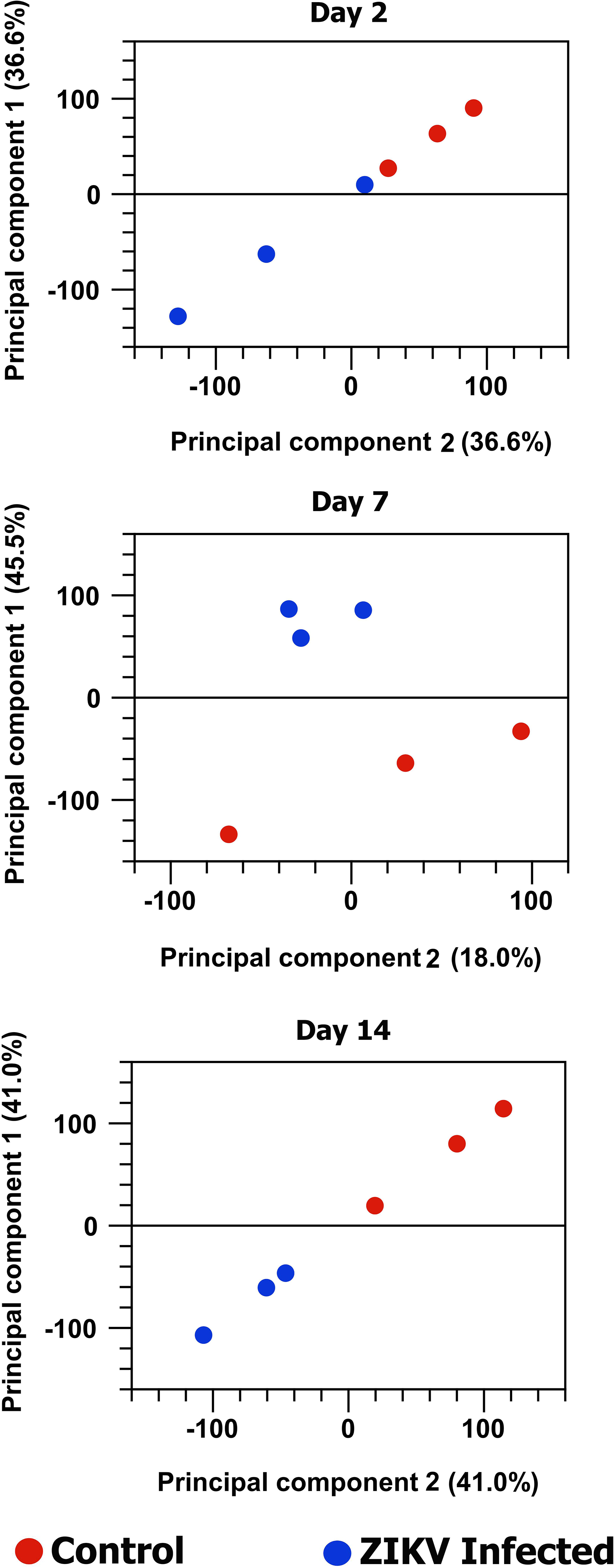
The principal component analysis of the effect of ZIKV infection on *Ae. aegypti* transcriptome at three different time points post-infection. The normalized log CPM (Count Per Million) used as expression value in this analysis.

Analysis and comparison of mRNA expression profiles of *Ae. aegypti* mosquitoes at different time points following ZIKV infection revealed that in total 1332 genes had changes of 2-fold or more in either directions (Fig. 2 with details in Table S2). Among the three time points, the highest number of changes occurred at 7 dpi with 944 genes showing alteration in their transcript levels. The numbers of genes altered at 2 and 14 dpi were very close, 298 and 303, respectively (Fig. 3). These trends were expected as we anticipated to see lower gene expression alteration at 2 dpi and 14 dpi due to low level of infection in the mosquito body at 2 dpi and advanced stages of virus replication at 14 dpi, while at 7 dpi the virus is still at its proliferative stage infecting various tissues of the mosquito. In a previous study that explored the effect of DENV-2 on *Ae. aegypti* transcriptome using RNA-Seq, the number of genes altered at 4 dpi was the highest (151 combining carcass and midgut) as compared to 1 dpi that showed the lowest number of changes (40) followed by 14 dpi (82) (11).

**Figure 2.**
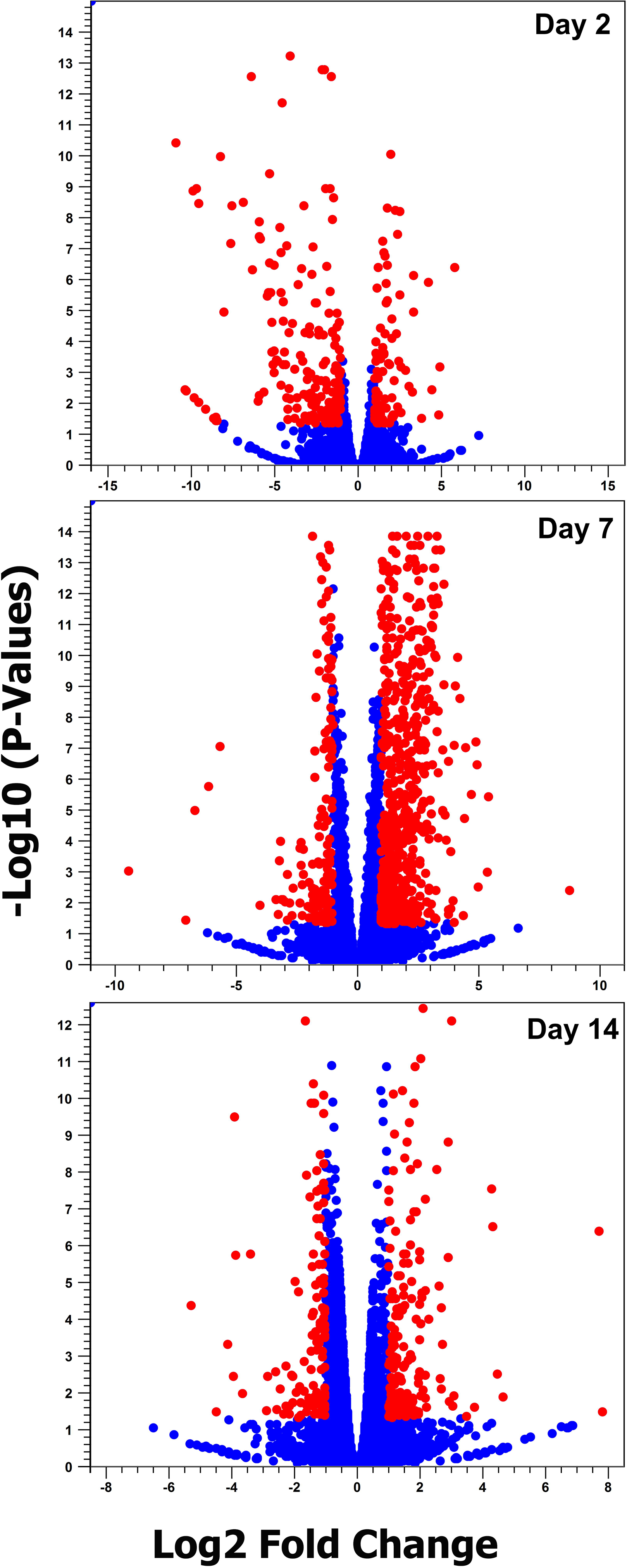
Volcano plot analysis. Red circles indicate differentially expressed mRNAs in response to ZIKV infection (Fold change > 2 and FDR < 0.05).

**Figure 3.**
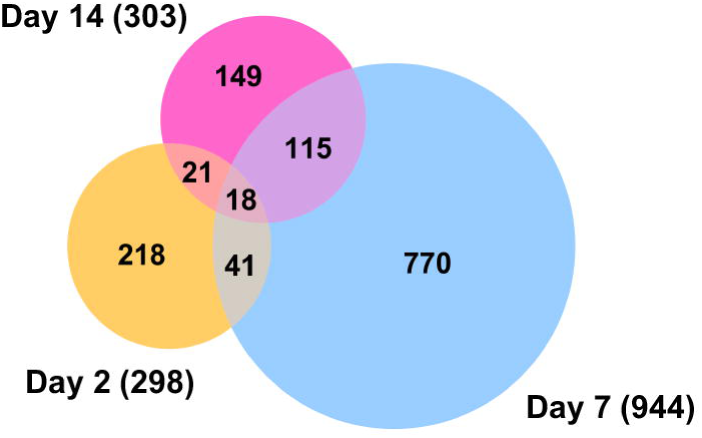
Venn diagram representing the number of differentially expressed coding genes at three different time points post ZIKV infection. Profound alteration in gene expression was observed at 7 dpi and more common differentially expressed genes were found between day 7 and 14 samples.

Comparison of the transcriptome profiles showed 18 overlapping genes among the three time points (Fig. 3; listed in Table 1). Twelve of these common genes were depleted and only six were enriched, which were Angiotensin-converting enzyme (AAEL009310), serine-type endopeptidase (AAEL001693), phosphoglycerate dehydrogenase (AAEL005336), cysteine dioxygenase (AAEL007416) and two hypothetical proteins. To validate the analysis of the RNA-Seq data, we used RT-qPCR analysis of the 18 genes. Overall, all expression values showed consistency between the two methods and had a positive linear correlation (Pearson correlation; Day 2 R^2^ = 0.7097 P<0.0001; Day 7 R^2^ = 0.8793 P<0.0001; Day 14 R^2^ =0.9184 P<0.0001) (Fig. 4).

**Table 1.**
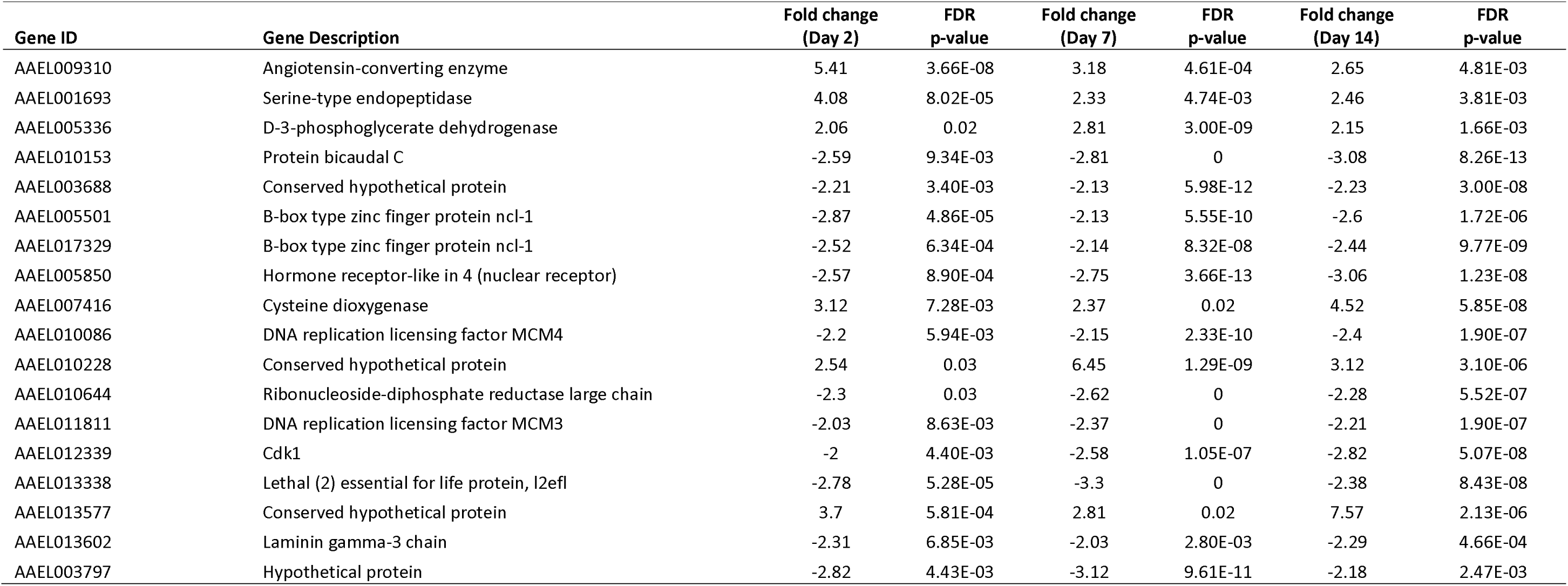
List of *Ae. aegypti* differentially expressed genes common to all the three time points post ZIKV infection.

**Figure 4.**
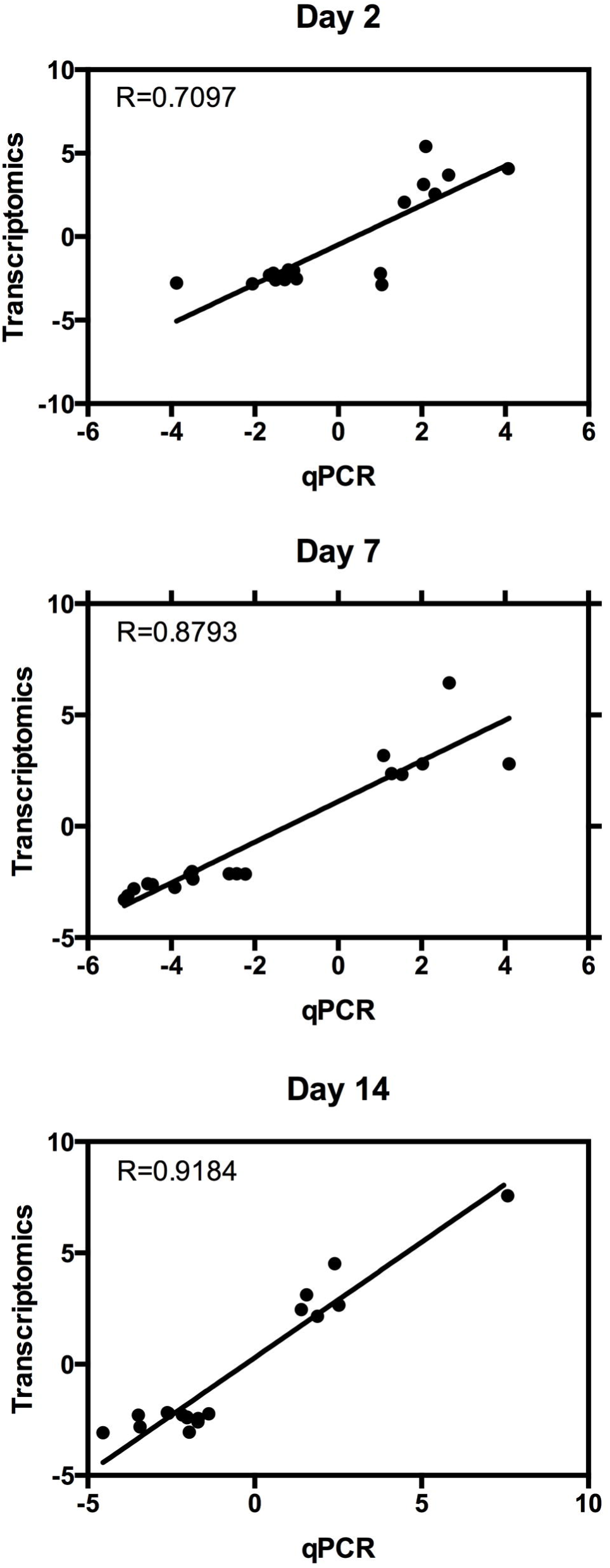
Validation of RNA-Seq data analysis by RT-qPCR. The 18 genes that were differentially expressed at all time points were validated by qPCR at 2, 7 and 14 days post-infection. Overall, all time points showed consistency between the two methods in their trends of depletion or enrichment.

### Differentially abundant transcripts and comparisons with other flaviviruses

When concentrating on genes with 10-fold differential expression and statistical significance relative to control mosquitoes, 101, 54 and 17 genes showed changes at 2, 7 and 14 dpi, respectively (Table S2 in dark blue font). After removing hypothetical proteins, those with known functions are listed in Table 2. Interestingly, while the total number of genes showing differential abundance was higher at 7 dpi (Fig. 3), more genes showed 10-fold or greater changes at 2 dpi as compared with 7 dpi (101 versus 54).

**Table 2.**
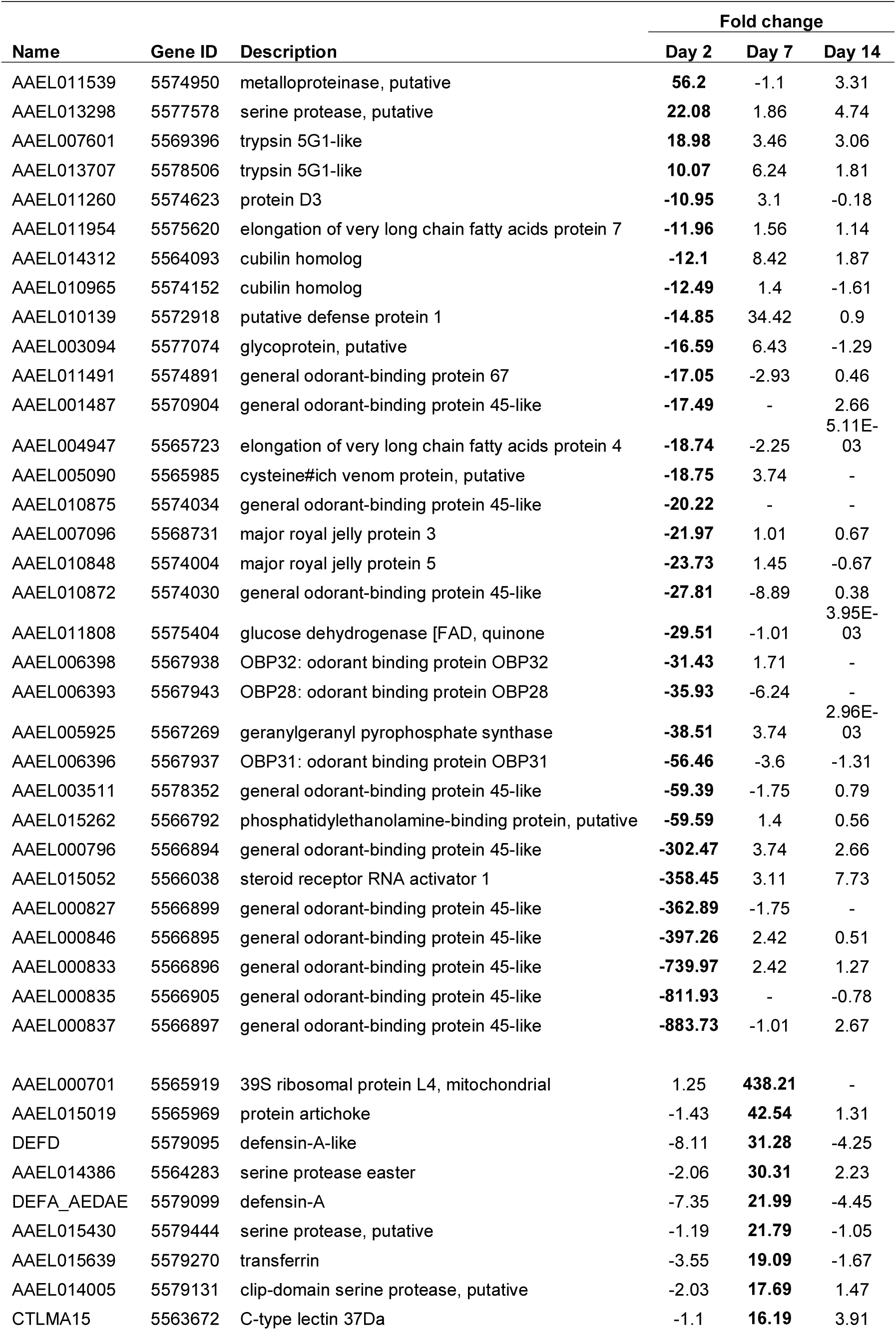
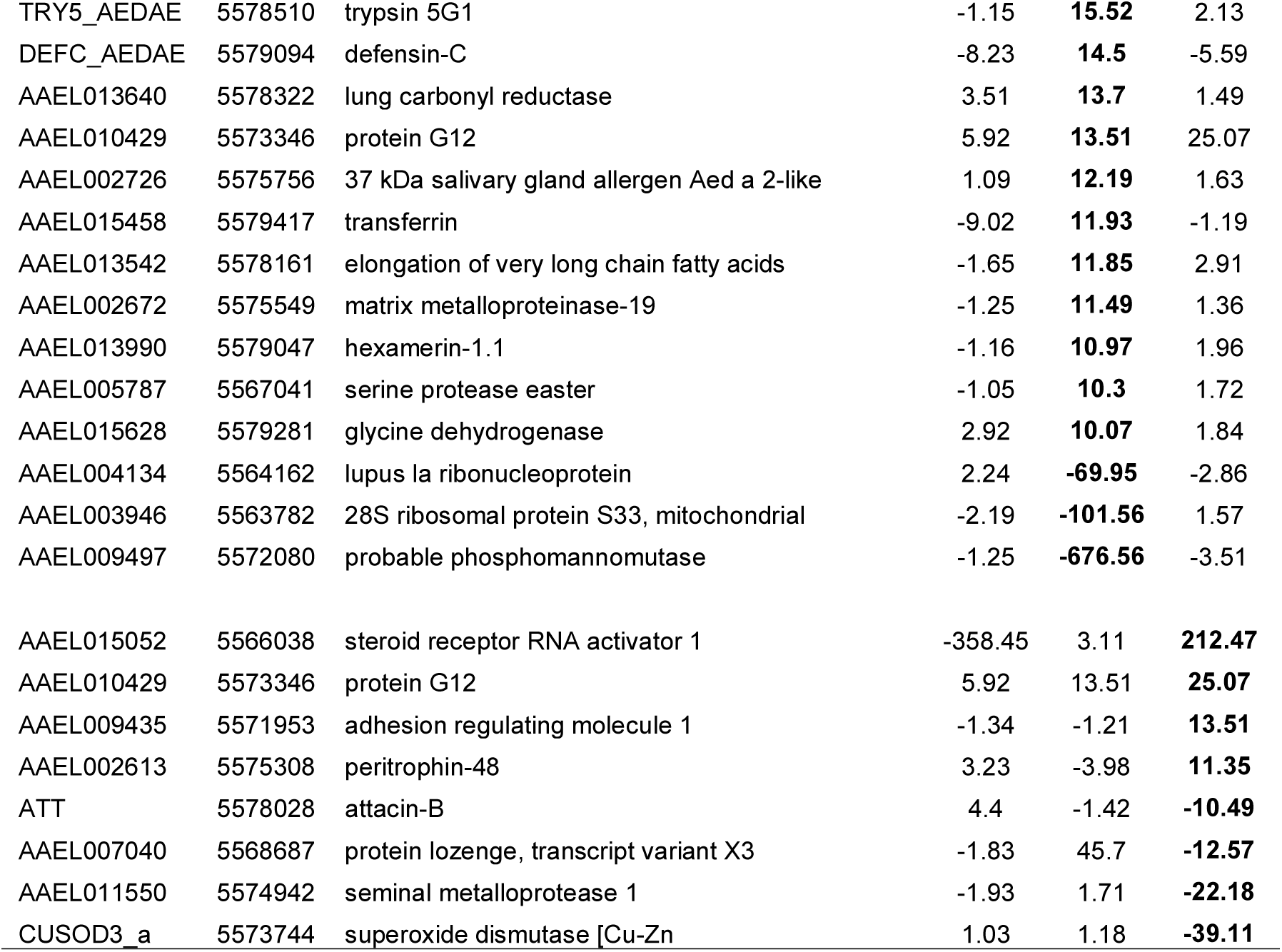
List of *Ae. aegypti* differentially expressed genes with more than 10-fold change specific to each time point (in bold) post ZIKV infection.

At 2 dpi, transcripts of eight genes were enriched with a metalloproteinase (AAEL011539) showing 56-fold increase in abundance, a serine protease (AAEL013298) increasing 22-fold, and two trypsins (AAEL007601 and AAEL013707) with 19 and 10-fold increases, respectively. We also saw that two phosphatidyl ethanolamine-binding proteins, two cubulin proteins, and a cysteine-rich venom protein were altered at this time point. However most strikingly, we observed suppression of 14 odorant binding proteins at 2 dpi with several of these transcripts being massively reduced (around 800-fold) (Table 2). Furthermore, other odorant binding protein transcripts were enhanced (by 2 fold or greater) at 7 and 14 dpi (Fig. S1), indicating that ZIKV may have the capacity to alter the behavior of the mosquito, potentially suppressing host-seeking in early stages of the infection and encouraging host-seeking when the mosquito is infectious. Dengue virus is known to alter host-seeking behaviors and feeding efficiency (17, 18), and microarray analysis of mosquitoes with salivary gland infections found several odorant binding protein transcripts that were enhanced in this late stage of infection (14 dpi) (19). Similarly, there is evidence that malaria parasites suppress the host-seeking tendencies of the mosquito early in infection but encourage host-seeking at later stages when the mosquito can transmit the parasite (20-22). The transcription patterns we observed here with ZIKV are consistent with these observations from dengue and malaria infection of mosquitoes but further behavioral studies are required to confirm this intriguing finding.

At 7 dpi, 34 genes showed enrichment of 10-fold or more including clip-domain serine proteases, defensins, transferrins, hexamerin, C-type lectin, and serine proteases, which are implicated in immune responses. At this time point, only seven genes were depleted. The number of genes that were differentially expressed by 10-fold or more at 14 dpi was small, with eight genes showing enrichment and eight genes showing depletion. The highest enrichment (212-fold) was steroid receptor RNA activator 1 (AAEL015052), while peritrophin, attacin and superoxide dismutase were among the depleted genes (Table 2). Previous studies have shown alteration of mRNA transcript levels in *Ae. aegypti* mosquitoes infected with DENV and a couple of other flaviviruses. Using microarray analysis, Colpitts et al. found that 76 genes showed 5-fold or more changes in DENV-infected mosquitoes over 1, 2 and 7 dpi (13). Their study, which also included response of *Ae. aegypti* to West Nile virus (WNV) and Yellow fever virus (YFV), found commonly 20 and 15 genes were differentially enriched and depleted, respectively, between the three flaviviruses at day 1 post-infection. Considering utilization of two different techniques in Colpitts et al (microarray) and in this study (RNA-Seq) and differences between the time points chosen, proper comparison of changes in transcript levels and fold changes cannot be done. However, when we mapped all the differentially expressed genes (2-fold or more) from Colpitts et al. against our data (Table S2), we found 364 genes from our study showed differential expression at least in one time point that overlap with the other three viral infections (Table S3).

In a follow-up study using the data from the above study (1, 2, 7 dpi) (13), Londono-Renteria et al. identified 20 top differentially regulated transcripts in YFV, DENV and WNV infected *Ae. aegypti* mosquitoes (23). Out of these 20 genes, five of them were also found changed in ZIKV-infected mosquitoes in our study. These were the cysteine-rich venom proteins (AAEL005098, AAEL005090, AAEL000379 and AAEL000356) by about 9, 18, 25 and 150-fold depletion at 2 dpi, and an unknown protein (AAEL013122) by 390-fold depletion at 2 dpi. While pairwise comparison is not really possible between the two studies, comparing data from 2 dpi showed that AAEL005090 (in the case of DENV), AAEL005098 and AAEL000356 (in the case of YFV and WNV), and AAEL013122 (in the case of DENV) changed in the same direction as ZIKV infection. Another study also found a number of cysteine-rich venom proteins altered upon DENV infection of *Ae. aegypti* mosquitoes (11). Cysteine-rich venom proteins are secretory proteins that are mostly found in the fluids of animal venoms acting on ion channels (24). Londono-Renteria et al. found that among the cysteine-rich venom proteins only AAEL000379 was enriched in DENV-infected mosquitoes and the rest did not change noticeably. Silencing the gene led to increase in replication of DENV (23). Alteration of the cysteine-rich venom proteins commonly found in the case of different flaviviruses indicates their possible importance in replication of these viruses. Further studies are required to determine the role these proteins play in ZIKV-infected mosquitoes specifically.

In another study with DENV-2 and *Ae. aegypti* in which deep sequencing of carcass, midgut and salivary glands with one replicate per pooled sample were used, transcript levels of infected and non-infected tissues were compared at 1, 4 and 14 dpi, which showed differential abundance of 397 genes (11). We reanalyzed the raw data from the study using the same pipeline as we used for our study. While comparative analysis of the study with ours cannot properly be made due to differences in the samples (tissues versus whole mosquitoes) and timings post-infection, in total, we found 199 genes commonly altered between DENV-2 and ZIKV infections, some with the same directional change in expression (Table S4).

A number of immune-related genes were mostly enriched at 7 dpi in ZIKV-infected mosquitoes. Toll was enriched only at 7 dpi by 2-fold. Twelve leucine-rich immune proteins were mostly enriched at 7 dpi by 4-16 folds. Phenoloxidae (AAEL010919), which was not changed upon DENV infection, was depleted by 2-folds at 2 dpi but enriched by 8-9 folds at 7 and 14 dpi in ZIKV-infected mosquitoes. Components of the JAK/STAT pathway, such as Dome and Hop, were not induced in ZIKV-infected mosquitoes. Interestingly, induction of the JAK/STAT pathway specifically in the fat body of *Ae. aegypti* mosquitoes by overexpressing Dome or Hop did not lead to increased resistance to ZIKV infection (25). This result and lack of induction of the pathway in our study suggests that the JAK/STAT pathway may not be involved in ZIKV-mosquito interaction. Further, major genes involved in the RNAi pathway, such as Dicer-1, Dicer-2, or any of the Argonaut genes, also did not change upon ZIKV infection in this study.

### Gene Ontology

All the differentially expressed host genes were submitted to Blast2Go for gene ontology (GO) analysis. This analysis identified 126, 68 and 33 GO terms in biological process, molecular function and cellular components, respectively (Table S5). GO analysis of enriched genes at different times post-infection showed that they were mostly related to proteolysis, zinc ion/protein binding and integral component of membrane (Fig. 5). Among the depleted genes, the highest categories were more variable, with day 2 having chitin metabolic process, odorant binding, integral component of membrane, at day 7 oxidation-reduction process, DNA binding and nucleosome, and at day 14 oxidation-reduction process, protein binding and nucleus (Fig. 5). In support of our earlier observation (Fig. S1), odorant binding transcripts were depleted at day 2 but enriched at day 14 (Fig. 5). In *Ae. aegypti*, differentially expressed genes upon infection with DENV, WNV and YFV belonged to various cellular processes, such as metabolic processes, ion binding, peptidase activity and transport (13), which are also among the GO terms identified in differentially abundant transcripts in the ZIKV-infected mosquitoes (Fig. 5). The genes commonly altered upon ZIKV and DENV infections listed in Table S4 were mostly in proteolysis, oxidation-reduction process and transmembane transport from the biological process, serine-type endopeptidase activity and protein binding from the molecular function, and integral component of membrane, nucleus and extracellular region from cellular component (Table S4).

**Figure 5.**
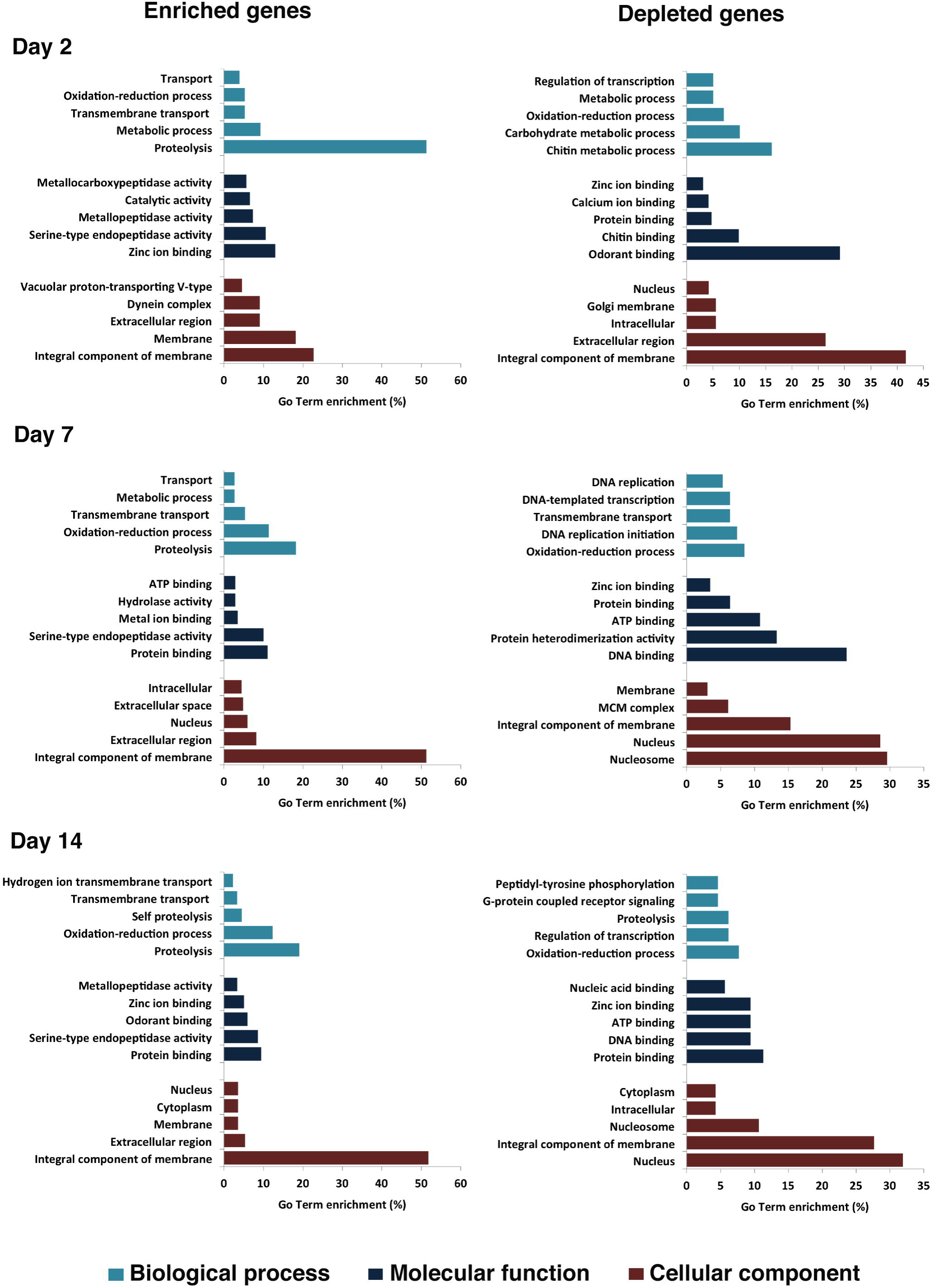
GO term enrichment analysis of differentially expressed genes in response to ZIKV infection in three categories of biological process, molecular function, and cellular component for enriched and depleted genes at 2, 7 and 14 days post-infection.

### microRNA target genes

Recently, we identified 17 *Ae. aegypti* microRNAs (miRNA) altered upon ZIKV infection at the same time points that RNA-Seq was conducted (2, 7 and 14 dpi) (16). Comparative analysis of the altered mRNAs and the 17 miRNAs with opposite trends in abundance revealed that 53 of the differentially expressed mRNAs could potentially be regulated by 11 out of the 17 differentially abundant miRNAs (Table S6). However, there is growing evidence that miRNAs could also positively regulate their target genes (26, 27), which are not listed in the table. Further, the analysis showed that some miRNAs have multiple potential target genes as expected (e.g. miR-309a has 19 target genes and miR-981-5p with 12 target genes). Gene ontology analysis of the target genes indicated that the majority of the genes are involved in oxidation-reduction process and integral component of membrane within the Biological Process and Cellular Component terms (Table S6).

### Long intergenic non-coding RNAs (lincRNAs) change upon ZIKV infection

lincRNAs are transcripts that are larger than 200 nt but do not code for any proteins, however, they are transcribed the same way as mRNAs (28); i.e. they have a poly-A tail and therefore enriched in transcriptomic data produced following mRNA isolation and sequencing. Similar to small non-coding RNAs, the main function of lincRNAs is regulation of gene expression, involved in various processes such as genomic imprinting and cell differentiation (29), epigenetic and non-epigenetic based gene regulation (30), activation and differentiation of immune cells (31), and relevantly virus-host interactions (32-36).

We recently reported 3,482 putative lincRNAs from *Ae. aegypti* (32). In this study, we found that in total, 486 lincRNAs were differentially expressed in response to ZIKV infection in at least one time point post-infection (fold change > 2 and P-value <0.05). Similar to mRNAs (see Fig. 3), the majority of altered lincRNAs were found at 7 dpi and 56 out of these lincRNAs showed significant alteration at least in two time points (Table S7 and Fig. 6). The Euclidean distance was calculated for each time point based on their lincRNA fold changes. Differentially expressed lincRNAs at 7 dpi (116.83) and 14 dpi (75.30) showed more correlation or similar fold-change pattern than those of 2 dpi (180.86). Only lincRNAs 656, 1385 and 3105 were differentially expressed and showed the same fold-change change pattern among the three time points. In our previous study, we also found that the transcript levels of 421 *Ae. aegypti* lincRNAs was altered due to DENV-2 infection. Comparison of those with the ones identified in this study showed that about 80 of them were also differentially expressed in ZIKV-infected samples (Table S7), which could be common lincRNAs involved in flavivirus-mosquito interactions.

**Figure 6.**
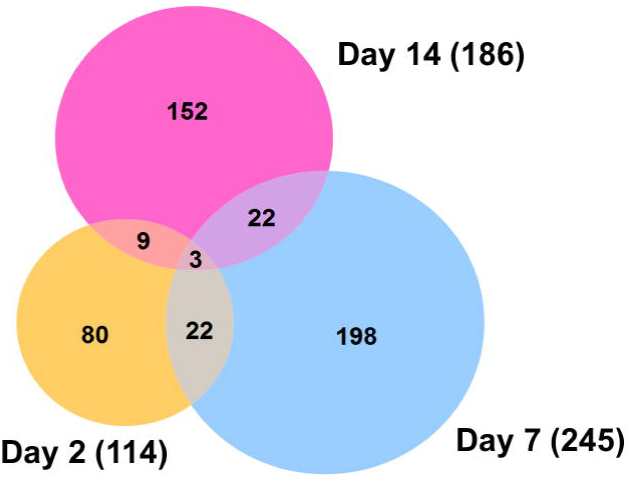
Venn diagram representing the number of differentially expressed lincRNAs at three different time points post ZIKV infection (fold-change > 2 and P-value <0.05). The majority of altered lincRNAs were found at 7 dpi and 56 out of these lincRNAs showed significant alteration at least at two time points.

## Conclusions

Overall, our results showed large changes in the transcriptome of *Ae. aegypti* mosquitoes upon ZIKV infection, both in coding and long non-coding RNAs. The majority of transcriptional changes occurred at 7 dpi, with the genes mostly involved in metabolic process, cellular process and proteolysis. We found some overlaps of transcriptional alterations in the case of other flavivirus infections in *Ae. aegypti*, but unlike those, immune genes were not altered to the same extent. In regards to lincRNAs, out of 486 lincRNAs changed in ZIKV-infected mosquitoes, 80 of them overlapped with that of DENV-infected mosquitoes indicating possible conserved functions of the lincRNAs in flavivirus-mosquito interactions. A drawback of this study is that we used whole mosquitoes, which means changes at the tissue levels could have been overlooked due to dilution factor by mixing all tissues; however, the outcomes provide a global overview of transcriptional response of *Ae. aegypti* to ZIKV infection, and can be utilized in determining potential pro and anti-viral host factors.

## Materials and Methods

### Ethics Statement

ZIKV, which was originally isolated from an *Ae. aegypti* mosquito (Chiapas State, Mexico), was obtained from the World Reference Center for Emerging Viruses and Arboviruses at the University of Texas Medical Branch (Galveston, TX, USA). Experimental work with the virus was approved by the University of Texas Medical Branch Institutional Biosafety Committee (Reference number: 2016055).

### Mosquito infections with Zika virus

We used excess RNA from samples generated recently to investigate miRNA profiles in ZIKV-infected *Ae. aegypti* mosquitoes (16). Briefly, 4-6 day old female *Ae. aegypti* (Galveston strain) were orally infected with ZIKV (Mex 1-7 strain) at 2 × 10^5^ focus forming units (FFU)/ml) in a sheep blood meal (Colorado Serum Company). Infected mosquitoes were collected at 2, 7 and 14 days post-infection (dpi) from which RNA was extracted using the mirVana RNA extraction kit (Life Technologies) applying the protocol for extraction of total RNA. Viral infection in mosquitoes was confirmed by Taqman qPCR on ABI StepOnePlus machine (Applied Biosystems) (16). For all time points, three independent pools were used to create libraries for infected and uninfected samples. Uninfected mosquitoes were fed with ZIKV-free blood, collected at the same time points and processed as above. The dynamics of infection in mosquitoes was shown in Saldaña et al. (16) in Fig. S1.

### Library preparations and sequencing

All samples were quantified using a Qubit fluorescent assay (Thermo Scientific). Total RNA quality was assessed using an RNA 6000 chip on an Agilent 2100 Bioanalyzer (Agilent Technologies).

Total RNA (1.0 μg) was poly A+ selected and fragmented using divalent cations and heat (94° C, 8 min). The NEBNext Ultra II RNA library kit (New England Biolabs) was used for RNA-Seq library construction. Fragmented poly A+ RNA samples were converted to cDNA by random primed synthesis using ProtoScript II reverse transcriptase (New England Biolabs). After second strand synthesis, the double-stranded DNAs were treated with T4 DNA polymerase, 5’ phosphorylated and then an adenine residue was added to the 3’ ends of the DNA. Adapters were then ligated to the ends of these target template DNAs. After ligation, the template DNAs were amplified (5-9 cycles) using primers specific to each of the non-complimentary sequences in the adapters. This created a library of DNA templates that have non-homologous 5’ and 3’ ends. A qPCR analysis was performed to determine the template concentration of each library. Reference standards cloned from a HeLa S3 RNA-Seq library were used in the qPCR analysis. Cluster formation was performed using 15.5-17 billion templates per lane using the Illumina cBot v3 system. Sequencing by synthesis, paired end 50 base reads, was performed on an Illumina HiSeq 1500 using a protocol recommended by the manufacturer.

### RNA-Seq data analysis

The CLC Genomics Workbench version 10.1.1 was used for bioinformatics analyses in this study. RNA-Seq analysis was done by mapping next-generation sequencing reads, distributing and counting the reads across genes and transcripts. The latest assembly of *Aedes aegypti* genome (GCF_000004015.4) was used as reference. All libraries were trimmed from sequencing primers and adapter sequences. Low quality reads (quality score below 0.05) and reads with more than 2 ambiguous nucleotides were discarded. Clean reads were subjected to RNA-Seq analysis toolbox for mapping reads to the reference genome with mismatch, insertion and deletion cost of 2, 3 and 3, respectively. Mapping was performed with stringent criteria and allowed a length fraction of 0.8 in mapping parameter, which encounter at least 80% of nucleotides in a read must be aligned to the reference genome. The minimum similarity between the aligned region of the read and the reference sequence was set at 80%.

Principal Component Analysis (PCA) graphs were produced for each time point after ZIKV infection between control and infected samples to identify any outlying samples for quality control. The expression levels used as input were normalized log CPM (Count Per Million) values.

The relative expression levels were produced as RPKM (Reads Per Kilobase of exon model per Million mapped reads) values, which take into account the relative size of the transcripts and only uses the mapped-read datasets to determine relative transcript abundance. To explore genes with differential expression profile between two samples, CLC Genomic Workbench uses multi-factorial statistics based on a negative binomial Generalized Linear Model (GLM). Each gene is modeled by a separate GLM and this approach allows us to fit curves to expression values without assuming that the error on the values is normally distributed. TMM (Trimmed mean of M values) normalization method was applied on all data sets to calculate effective library sizes, which were then used as part of the per-sample normalization (37).

The Wald Test was also used to compare each sample against its control group to test whether a given coefficient is non-zero. We considered genes with more than 2-fold change and false discovery rate (FDR) of less than 0.05 as statistically significantly modulated genes.

We previously reported 3,482 putative long intergenic non-coding RNAs (lincRNA) from *Ae. aegypti* using a stringent filtering pipeline to remove transcripts that may potentially encode proteins (32). The expression profile of lincRNAs was also generated for each sample similar to the approach described above.

To identify the host transcriptomic response to two different flaviviruses, we compared altered gene profiles in previously published DENV-infected *Ae. aegypti* libraries (11) with our ZIKV infected samples. The relevant RNA-Seq data (SRA058076) were downloaded from NCBI sequence read archive. The libraries were treated in the same way as described above to identify differentially expressed *Ae. aegypti* gene profiles in response to DENV.

### Gene Ontology (GO) analysis

All differentially expressed genes were uploaded to Blast2GO server for functional annotation and GO analysis. We used Blast and InterProScan algorithms to reveal the GO terms of differentially expressed sequences. More abundant terms were computed for each category of molecular function, biological process and cellular components. Blast2GO has integrated the FatiGO package for statistical assessment and this package uses the Fisher’s Exact Test.

### Identification of miRNA target genes

We screened all differentially expressed mRNAs to identify potential miRNA targets among them. If selected mRNAs do not have complete annotation such as clear 5’UTR, ORF and 3’UTR, the region before ORF start codon (300 bp) and after stop codon (500 bp) for each mRNA was considered as 5’UTR and 3’UTR, respectively. We used three different algorithms including RNA22 (38), miRanda (39) and RNAhybrid (40) to predict potential miRNA binding sites on genes altered by ZIKV. We previously described this approach and parameters for setting each tool, but to increase the level of confidence, we selected those binding sites which were predicted by all the three algorithms for further analysis (41).

### RT-qPCR analysis of mRNAs

qPCR validations were done using the same RNA that was used for RNA-Seq. RNA from ZIKV positive samples was pooled (N = 5) for time points 2, 7, and 14 dpi and treated with amplification grade DNase I (Invitrogen). Total RNA was reverse transcribed using the amfiRivert cDNA Synthesis Platinum master mix (GenDEPOT, Barker, TX, USA) containing a mixture of oligo dT(_18_) and random hexamers. Real-time quantification was performed in StepOnePlus instrument (Applied Biosystems, Foster City, California, United States) in 10 µl reaction containing 1:10 diluted cDNA template, 1X PowerUp SYBR Green Master Mix (Applied Biosystems), 1µM each primer. The analysis was performed using ΔΔCt (Livak) method (42). Three independent biological replicates were conducted and all PCRs were performed in duplicates. The ribosomal protein S7 gene (43) was used for normalization of cDNA templates. Primer sequences are listed in Table S8.

### Accession number

The accession number for the raw and trimmed sequencing data reported here is GEO: GSE102939.

## Acknowledgements

This project was supported by a NIH grants (R21AI124452 and R21AI129507) and a University of Texas Rising Star award to GLH, and an Australian Research Council (DP150101782) and an Australian Infectious Disease Research Centre grant to SA. GLH is additionally supported by the Western Gulf Center of Excellence for Vector-borne Diseases (CDC grant CK17-005). MAS was supported by a NIH T32 fellowship (2T32AI007526) while SH was supported by a James W. McLaughlin postdoctoral fellowship at the University of Texas Medical Branch. We would like to thank the University of Texas Medical Branch insectary core for providing mosquitoes and the World Reference Center for Emerging Viruses and Arboviruses for providing the ZIKV isolate.

## Author contributions

conceptualization, S.A., and G.L.H; Investigation: K.E., S.H., M.A.S., S.G.W., and T.G.W.; Data curation: K.E.; Formal analysis: K.E., and S.A.; Writing-original draft: K.E., and S.A.; Writing-review & editing: S.A., and G.L.H.; Supervision: S.A., and G.L.H.

## Supplemental material legends

**Figure S1**. Expression levels of odorant binding protein transcripts at days 2, 7, and 14.

**Table S1**. RNA read summary in ZIKV-infected and non-infected libraries.

**Table S2**. Differentially expressed transcripts in response to ZIKV at 2, 7, and 14 dpi.

**Table S3**. Comparison of transcripts modulated by ZIKV to those modulated by dengue virus, West Nile virus and Yellow fever virus identified by Colpitts *et al.* (13).

**Table S4**. Comparison of transcripts modulated by ZIKV to those modulated by dengue virus identified by Bonizzoni *et al.* (11).

**Table S5**. Gene ontology analysis for transcripts differentially regulated by ZIKV.

**Table S6**. ZIKV regulated transcripts potentially affected by miRNAs identified by Saldaña *et al*. (16).

**Table S7**. Differentially expressed lincRNAs in response to ZIKV at 2, 7, and 14 dpi.

**Table S8**. List of primers used in the study.

